# Force, angle, and velocity parameters of finger movements are reflected in corticospinal excitability

**DOI:** 10.1101/2024.02.28.582459

**Authors:** I.M. Brandt, J. Lundbye-Jensen, T. Grünbaum, M.S. Christensen

## Abstract

Identifying which movement parameters are reflected in the corticospinal excitability (CSE) will improve our understanding human motor control. Change in CSE measured with transcranial magnetic stimulation (TMS)-induced motor evoked potentials (MEPs) can probe the content of the signal from primary motor cortex (M1) through the corticospinal pathway and spinal motoneurons to the muscle. Here we used MEPs to investigate which movement-related parameters are reflected in CSE in 33 healthy adults. In three separate tasks, we evaluated which movement parameter(s), force, angle, and velocity, are reflected in the MEP amplitude during movement preparation and movement execution. Bayesian model comparison in a forward feature selection framework identified force and velocity measures as reflected in the MEP amplitude during movement preparation, and force, angle, and velocity measures as reflected in the MEP amplitude during movement execution. Importantly, we included measures of electromyography (EMG) in the forward feature selection, and the parameter measures are included only if they add explanatory power of MEP amplitude in addition to the EMG. These findings show that when taking EMG measures into account, all three movement parameters force, angle, and velocity are reflected in CSE. These findings propose a flexible and task-dependent form of signaling in the motor system that allows parameter-specific modulation of CSE to accurately control finger movements.

**Key points:**

- Prior research show that the primary motor cortex activity reflects movement parameters.
- Measures of the response to a magnetic stimulation, the motor evoked potential (MEP), can be used to assess the content of the signal sent to the muscle.
- We use Bayesian model comparison to test whether movement parameters are reflected in the models best describing the MEP amplitude modulations.
- We show that the MEP amplitude reflects all tested movement parameters, force, angle, and velocity.
- Our results indicate a task-dependent form of signaling not only in M1, but also in the corticospinal pathway and spinal motor neurons propagating the signal to the muscle.

## Introduction

Understanding human motor control requires identifying which parameters of movement, e.g. contraction force, angle, and movement velocity, are reflected in the motor system. The primary motor cortex (M1) and the corticospinal pathway have central roles in coordination and execution of movements. Studies in primates show that M1 neuronal activity is correlated with load direction/force (Kalaska et al., 1989), movement direction (Georgopoulos et al., 1986; Georgopoulos and Carpenter, 2015), velocity (Reina et al., 2001; Stark et al., 2007), and end position (Aflalo and Graziano, 2006). Stark et al. (2007) reported that 2/3 of motor neurons predominantly code for one movement parameter, most neurons representing velocity (Stark et al., 2007). These findings suggest that primate M1 neuronal activity is tuned to movement parameters in a complex, multidimensional manner (Aflalo and Graziano, 2006).

Motor commands are believed to originate from M1, propagate through the corticospinal pathway (CSP) and lead to activation of spinal cord motoneurons which activate target muscles. Cortical facilitation and disinhibition activate cortical output neurons, and these mechanisms increase corticospinal excitability (CSE). To probe changes in CSE, transcranial magnetic stimulation (TMS) can be applied to M1 to activate the cortex and induce motor-evoked potentials (MEPs) in muscles – primarily - of the contralateral limb. The MEP amplitude is modulated by state changes in the motor system such as movement planning and execution and can therefore be used as a read-out of changes in the motor system before and during a movement (Bestmann and Krakauer, 2015; Day et al., 1987).

Previous research has investigated the relationship between MEP amplitudes and movement parameters in human. MEP amplitude is tuned to wrist movement direction (Kadota et al., 2014) and shoulder position (Ginanneschi et al., 2005). MEP amplitude changes with force but not in a linear fashion (Kiers et al., 1995). Sasaki et al. (2018) found that during passive movements, velocity, and joint angles of the index finger movement increased MEP amplitude (Sasaki et al., 2018). Taken together, while MEP amplitude correspond to different movement parameters, some parameters have been assessed only during passive movements. It can therefore be speculated whether MEPs during passive movement to a larger extend represent modulations related to sensory feedback than movement related excitability changes (Sasaki et al., 2018). Consequently, it is important to investigate if MEP amplitudes prior to and during motor output reflect a range of movement parameters, like the primate M1 does during voluntary movement (Georgopoulos et al., 1986; Georgopoulos and Carpenter, 2015; Kalaska et al., 1989). This would involve investigating whether objective measures of force, angle, and velocity are reflected in the MEP amplitude during movement preparation and execution. The aim of this study was to investigate which movement parameter(s) the MEP amplitude reflects, to better understand the importance of force, angle, and velocity parameters in the coding of signals involved in motor control. Our experimental design comprises a task for each of the three parameters force, angle, and velocity, with TMS stimulation during either movement preparation or movement execution for each trial. The relation between MEP amplitude modulations and the three parameters are tested in distinct tasks, to test the effect of each parameter separately. In this paper the movement parameter ‘angle’ is used interchangeably with how the movement parameter ‘position’ is used in the motor control literature (Norup et al., 2023), as the movement is performed as planar flexion-extension of a single joint. Participants aimed for predetermined parameter levels, e.g. a force level 5 that corresponded to 50% of maximal voluntary contraction (MVC) force. This was performed without augmented or visual feedback of their own movement, to avoid the potential influence of visual feedback on MEP amplitudes (Bell et al., 2018; Senna et al., 2015). We employ Bayesian model comparison to explore which movement parameters best explain the amplitude of the MEP normalized to EMG during MVC. We build linear Bayesian models with the dependent variable MEP and relevant movement parameters and EMG measures as independent variables to test the effect of the independent variables with Bayesian model comparison in a forward feature selection framework.

Delineating which movement-related parameters that contribute with significant variance to CSE would improve our understanding of how the motor system communicates and which type of information is conveyed to the muscles.

## Experimental procedures

### Participants

Out of 40 included participants, 33 healthy volunteers (21 female, mean age 29.12, age range 20-45) completed the study (2 failed to complete all three tasks, 3 were excluded due to too high coil-to head distance, 1 terminated the experiment, and for 1 participant, TMS was activating her migraine hotspot). Participants were recruited through forsøgsperson.dk and University of Copenhagen (UCPH). TMS was performed in accordance with TMS safety guidelines (Rossi et al., 2021). A safety questionnaire developed from Rossi (2009) (Rossi et al., 2009), was used to exclude participants with contraindication to TMS safety. Prior to experiments, we obtained written informed consent from all participants for participating in the study and for collection and processing of their data in accordance with EU general data protection regulation. This study conformed to the Declaration of Helsinki (“World Medical Association declaration of Helsinki: Ethical principles for medical research involving human subjects,” 2013) and the protocol was approved by the ethics committee of The Capital Region of Denmark (H-21061035). Data collection and storage was performed in accordance with the guidelines approved by UCPH (2004334 - 4242). Participants were compensated with 150 DKK per hour for participating in the experiment.

### Questionnaires

Thirty-one participants were right-handed and two were ambidextrous when assessed with the Edinburgh handedness inventory (Oldfield, 1971) (83 +/-22 (mean +/- 1 sd)). Level of physical activity was assessed using International Physical Activity Questionnaire (IPAQ) (Craig et al., 2003), experience with fine motor skill tasks and hours of sleep prior to the examination day was assessed using a self-developed questionnaire.

### Experimental Design

In order to study which movement parameters are reflected in the CSE, we examined whether force, angle, and/or velocity are reflected in the TMS-induced MEP in three right index finger movement tasks: a force task, an angle task, and a velocity task.

Each task consists of trials in which participants aim for a specific level of the movement parameter (e.g. force level 5), indicated by tone pitch. The level is relative to the individual maximal effort in the respective task, as described below. For each trial an MEP is evoked by TMS either before or during the movement. We use *movement preparation* and *movement execution* throughout the manuscript to indicate whether the TMS pulse is applied during movement preparation or during movement execution. For all tasks, the inter-tone interval was randomized and for force and angle the participants were instructed to react as fast as possible. This ensured for force and angle preparation that the stimulation was timed to the movement preparation phase. Taken together, for each trial we obtained an MEP amplitude and an objective measure of the parameter, e.g. the maximum force in each force trial.

Task order was randomized between participants, with 5-7 consecutive blocks of 6 min of each task, the last block was always a sham condition. Given the substantial influence of vision on the motor system in facilitating motor control and possibly affecting the CSE, we prevented visual feedback. Participants therefore relied only on proprioception, somatosensation and motor command signals for controlling their contractions and movements.

We further investigated the effect of attentional focus (AF) on MEP amplitude. Participants were instructed to either pay attention to the feeling of the movement parameter (force, angle, velocity) in their finger (internal AF) or to pay attention to imagined visual external feedback (external AF). Experiments comparing motor performance during internal versus external attentional focus usually gives an unfair advantage to the external attention condition, since participants use visual information to a higher degree in this condition (Lohse et al., 2014). To circumvent this, the external attention condition in this experiment relies on the same informational content as internal attention but the participant is instructed to represents their performance as its visual consequence on a thermometer.

#### Preparations

We tested each participant during two 3-3.5 hour sessions separated by a long lunch-break in a lab visit over one day (two days for one participant). We obtained informed consents and conducted the TMS safety questionnaire and other questionnaires. We placed electromyography (EMG) electrodes over relevant right forearm and hand muscles (see below) and measured maximal voluntary contraction (MVC) of the right hand index finger. Right hand index finger flexion MVC was measured with the forearm in a semiprone position, palm facing medially and index finger straight. Participants pressed maximally for three seconds, waited 90 seconds, and again pressed maximally for three seconds. MVC was identified as the maximum force level measured in either of the two presses. We measured the maximal compound muscle potential (Mmax) from flexor digitorum profundus (FDP) (see below) and determined TMS stimulation hotspot and motor threshold (MT). Participants were seated comfortably with the right forearm resting horizontally on a table. The forearm was placed in a semiprone position, with the palm facing medially. Their arm and hand were covered by a cloth (force) or a box (angle, velocity) to prevent visual feedback. Breaks were given when needed.

#### Force task

The force task was to press isometrically with the right index finger against a stationary force transducer in a horizontal direction (Fig. 1). All fingers pointed straight ahead, and the arm position was stabilized by two sticks against which the participants could lean their arm. Participants were instructed to press at either level 1, 3, or 5 on a scale from 1-10, where 10 was their MVC and level 1, 3, and 5 were relative to MVC (10%, 30%, and 50%). Force levels were not higher than 5 to avoid fatigue. After measuring MVC, the force levels 1, 3, 5 were guided by verbal feedback 2-3 times. After initial guidance and indication of force levels, the participant did not receive any external feedback throughout the experiment. Participants had to rely on their own sensorimotor experience. For each trial, two auditory tones were played. Tone pitch level of the first tone indicated which force level the participant should aim for. Low, middle, and high pitch tone indicated level 1, 3, and 5, respectively. The second tone was a ‘go-tone’, played after a randomized inter-tone interval (1-2 s). Mean single-stimulus inter-stimulus interval in the force task was 6.0 s. Each participant performed 5-7 blocks of 6 minutes (last block was sham) with a mean of 137 force movement preparation trials (TMS stimulation during preparation), 90 force movement execution trials (TMS stimulation during movement) and 59 Sham trials per participant (TMS coil positioned with one wing perpendicular to participant’s head). For internal AF, participants were asked to ‘*pay attention to the feeling of the force in the finger’*, for external AF, participants were asked to ‘*imagine a thermometer, that gives you feedback on the force you are producing’*.

**Figure 1.**
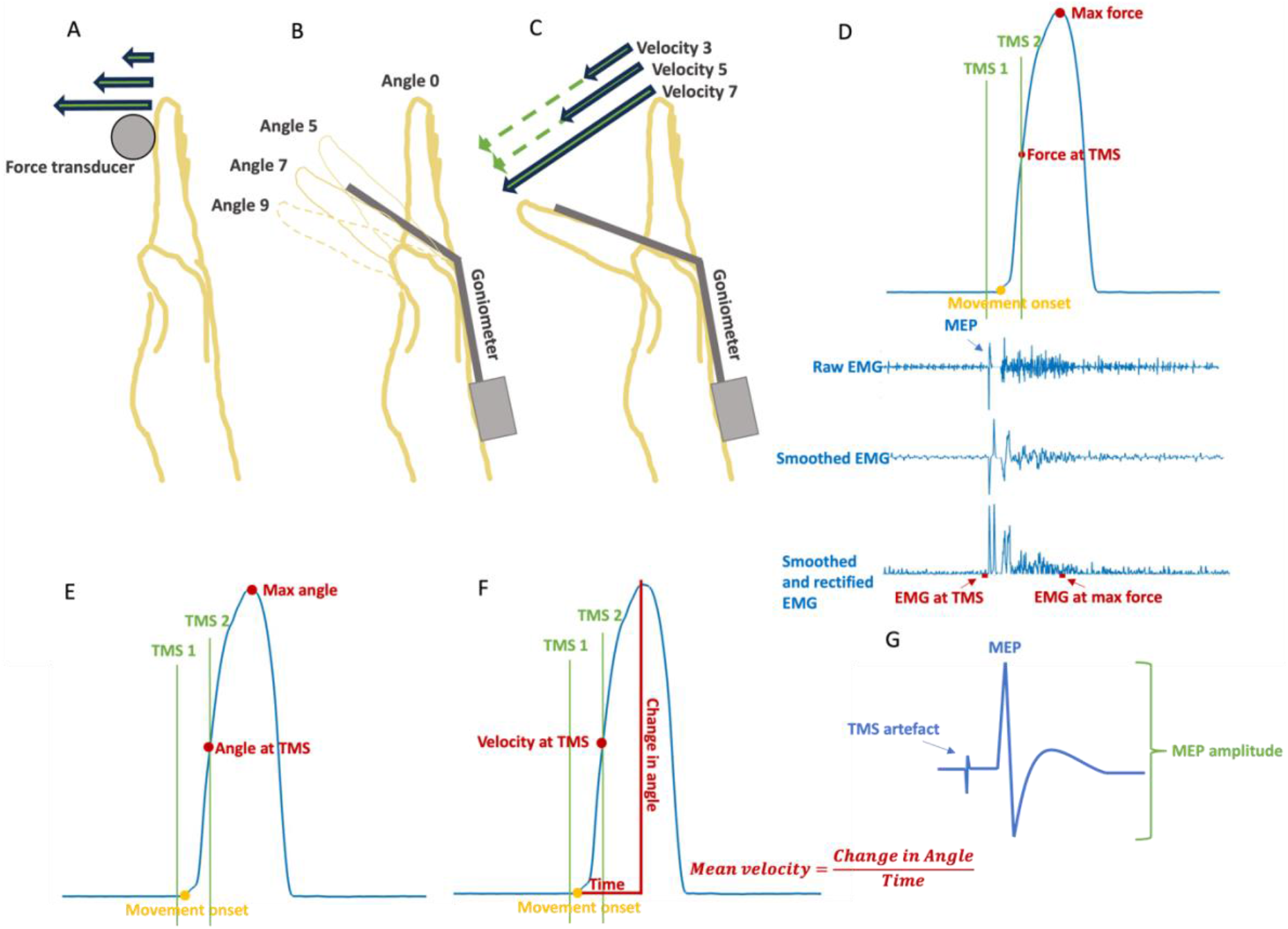
Experimental setup and tasks. A) Force task, right index finger was pressing isometrically on force transducer. B) Angle task, right index finger movements from angle 0 to angle 5 or angle 7. Angle 9 is shown to indicate the scale range. Movements were measured with goniometer. C) Velocity task, right index finger movements from start position to end position with velocity 3 (slow), 5, or 7 (fast). Movements were measured with goniometer. D) Example of force data with variables. Yellow circle indicates movement onset. Red circles indicate *max force* and *force at TMS*. TMS 1 indicates TMS stimulation during movement preparation, TMS 2 indicates TMS stimulation during movement execution. Raw, smoothed, and smoothed and rectified EMG data is shown below. In this example the TMS stimulation was applied during movement preparation, inducing a pre-movement MEP. E) Example of angle data with variables. F) Example of velocity data with variables. Mean velocity was calculated as change in angle from movement onset to max angle during the trial. Velocity at TMS was calculated as the mean velocity of smoothed and normalized data over 25ms prior to TMS stimulation to 25ms post TMS stimulation. G) Shows a drawing of MEP and the amplitude of the MEP. A TMS artefact is seen just prior to the MEP.

#### Angle task

The Right arm and hand were positioned as in the force task but with the force transducer removed. Right index finger movements were recorded with a fiber optic goniometer. Index finger pointing straight ahead was instructed as the starting point, angle 0. Participants were guided to learn to return to the same starting point. Max metacarpophalangeal (MCP) joint flexion was instructed as angle 9 (no bending in proximal interphalangeal joint was allowed). On this scale, participants were instructed to move to level 5 or 7 in each trial. Level 5 and 7 were guided 2-3 times by verbal feedback before initiation of task, and no more external feedback was provided. As in the force task, for each trial two tones were played. Angle level was indicated by low or high pitch tone, followed by a go-tone after a randomized inter-tone interval (1-1.5 s). Each participant performed 5-7 blocks of 6 minutes (last block was sham condition) with a total a mean of 67 angle movement preparation trials, 272 angle movement execution trials, and 66 sham trials per participant. Mean single-stimulus inter-stimulus interval during the experiment was 5.3s.

#### Velocity task

The velocity task resembled the angle task in position and movement and was likewise recorded with a goniometer. Participants were instructed to move right index finger from a position of pointing the finger straight ahead to max MCP joint flexion with different velocities. A low, middle, and high pitch tone indicated velocity level 3, 5, or 7, respectively, on a scale where velocity 1 was approximately 2 seconds, and velocity 9 was as fast as possible. Velocity levels were guided 2-3 times by verbal feedback before initiation of task, and no more external feedback was provided. The level-indicating tone was followed by a go-tone after a randomized inter-tone interval (1-2 s). Each participant performed 5-7 blocks of 6 minutes (last block was sham condition) with a mean of 82 velocity preparation trials, 232 velocity movement trials and 63 sham trials per participant. Mean single-stimulus inter-stimulus interval during the experiment was 6.2s.

The force task involved isometric contractions whereas the angle and velocity tasks involved concentric contractions and hence dynamic movements.

### Data acquisition

Experiments were run by a 1401 Micro Mk II analog-to-digital (AD) converter (Cambridge Electronic Design (CED), Cambridge, UK) and controlled by the software Spike2 (version 7.10, CED, Cambridge, UK). Spike2 controlled the signals for the TMS and sound through the CED, and signals from EMG, fiber optic goniometer (S720, Measurand Inc., New Brunswick, Canada), and force-transducer (UU2 load cell, DN-AM310 amplifier, Dacell, Seoul, Korea) were sampled at 2000 Hz with the CED. The EMG signal was amplified x20-100 (corrected for in the analysis) by a custom-build differential amplifier (Panum Institute electronics lab, Copenhagen, DK) and band-pass filtered (5-1000Hz). All data was stored on a safe drive.

Surface EMG was recorded from FDP muscle, first dorsal interosseous muscle (FDI) and extensor digitorum communis (EDC) using Ag/AgCl surface electrodes (dimensions 22*30 mm, Ambu A/S BlueSensor N-10-A/25 ECG electrodes, Ballerup, Denmark) in a belly-tendon montage. The active and the reference electrode were placed next to each other on the belly of the muscle, and the ground electrode was placed on the olecranon of the right ulnar bone. We used electrode gel to increase conductance (Signa Gel, Parker Laboratories, Inc., Fairfield, New Jersey 07004, USA). Participants skin was prepared with fine sandpaper and sanitizer (Hupfeld et al., 2020). Mmax was elicited by electric stimulation of the median nerve proximal to the elbow (pulse width: 200 μs, amplitude: 400 V, variable electrical current up to 100 mA) and recorded from the FDP muscle (stimulator model DS7A, Digitimer Ltd, Welwyn garden city, England). Although we recorded Mmax, we chose to normalize EMG data to EMG during MVC, since this better represents voluntary activation (Diong et al., 2022), and because the Mmax measurement was not reliable in some participants.

#### TMS stimulation and MEP measurements

Participants were stimulated with TMS over hand-M1 once per trial. Magnetic single-pulse stimulations were applied with a Magstim Rapid2 stimulator (Rapid2, The Magstim Company Ltd., Whitland, UK), using a custom-made batwing figure-of-8 shaped coil (biphasic pulse, initial Anterior (A)-to-Posterior (P) direction followed by PA direction, external coil diameter 115 mm). The coil was positioned horizontally over M1 with the handle of the coil pointing posterolaterally in a 45° angle to the coronal and/or sagittal plane. Resting MT (rMT) was determined by gradually increasing intensity of stimulator output until TMS induced MEPs (>50μV) in 5 out of 10 trials with an 8 second inter-stimulus interval (Hupfeld et al., 2020). Stimulation intensity in the experimental trials was 120% rMT. FDP motor hotspot was determined by finding a reactive spot lateral to the medial fissure and testing MEPs anterior, posterior, medial, and lateral to the measured starting point. Hotspot was the location with the largest and most consistent MEPs measured in the FDP muscle. A BrainSight® neuronavigational system (Rogue Research Inc, Montreal, Canada) was used to ensure consistent coil position and thus stimulation of the hotspot (below ∼3 millimeters from the identified hotspot). The coil was held by a mechanical coil-holder, and the head position adjusted if movement resulted in misalignment with the hotspot. In the sham condition, the coil was positioned with one wing perpendicular to participant’s head.

### Data analysis and statistics

#### EMG analysis

The raw EMG data was corrected for amplification (x20-100), zero-mean centered, smoothed over 40 ms, normalized to EMG at MVC, and rectified. MEP amplitude was determined from zero-mean centered raw EMG data and afterwards normalized to max EMG during MVC. MEPs with an amplitude less than three standard deviations (sd) from background EMG were excluded from the analysis. Although we have EMG measurements from FDP, FDI, and EDC muscles, we chose to conduct the analysis on FDP MEPs only. This decision was made to simplify the analysis process, acknowledging that the outcomes may vary among the muscles responsible for different functions during the movement. EMG at time of TMS is mean smoothed over 40 ms, normalized, and rectified EMG 55-5 ms prior to TMS stimulation. EMG at parameter measure is mean smoothed over 40 ms, normalized and rectified EMG 50-0 ms prior to max parameter measure.

#### Behavioral statistical analysis

We defined Bayesian linear mixed models of the parameter level participants were aiming for (e.g, force level 1, 3, or 5) explaining parameter measures, i.e. max force, max angle, mean velocity. The models were implemented in the library ‘brms’ (Bürkner, 2017) in the statistical software program R (https://www.r-project.org) using STAN (Carpenter et al., 2017). Subsequently, we performed Bayesian hypothesis tests to evaluate whether participants can distinguish between parameter levels. Width of credibility interval (CI) was used to evaluate the precision for each parameter level. Movement onset was defined as the time at which 5% of max angle in the trial was reached for angle and velocity, and for force it was defined as the time at which 2% of the max force in the trial was reached. Reaction time was measured as time from second tone onset to movement onset.

#### Determining best model explaining FDP MEP amplitude using Bayesian model selection

The data analysis was performed in MATLAB (version 9.13 (R2022b), The MathWorks Inc, Natick, MA, USA) and statistical analysis was performed in the program R (version 2022.12.0, https://www.r-project.org). For each movement parameter, force, angle, and velocity, we determined the model best explaining the CSE, measured as the FDP MEP amplitude, for both movement preparation and movement execution, resulting in 6 winning models in total. We used Bayesian tests in a Forward Feature Selection framework to compare the relative strength of competing models. We used the ‘brms’ package (version 2.18.0) in R to fit Bayesian linear mixed models and estimate model parameters. Models were compared using Bayes factors, which enables one to estimate the best model by computing bayes factors (BF) when comparing models to select the model with different independent variables using the ‘brms’ package bayes_factor function in R. We gathered samples with Markov Chain Monte Carlo sampling with 4000 iterations, 2000 warm-ups, and 7 chains, leading to 14.000 posterior samples. For each model run with brms, chain convergence was assessed with the Rhat statistic and a value of 1.00 was achieved.

Forward Feature Selection is an iterative method. The simple model included subject as a random effect with intercept, and in each iteration, we added the variable which improved the model the most, until no variable added further explanatory power (i.e., BF > 3).

We tested the variable *parameter measure*, which is *max force* or *max angle* or *mean velocity* during each trial, depending on task (Figure 1). We used *mean velocity* instead of *max velocity* since the focus of the task was the velocity from start to end of the movement. *Parameter measure at TMS* (force, angle, or velocity at time of TMS stimulation) was included for the respective parameters in the movement execution models. *Parameter level*, the level of the parameter participants are aiming for as explained in *experimental design,* was also tested, as well as an interaction factor between parameter measure and parameter level. *Attentional focus (AF)* is internal AF or external AF, also explained in *experimental design*. *EMG at TMS* and *EMG at max parameter* was also tested as variables. We further tested *block* as a variable to account for a possible change in MEP amplitude over time and included subject in all models. In the movement preparation models, *time from TMS to movement onset* was tested. We defined the variable *time from TMS to movement onset* as whichever of two events occurs first: an EMG increases exceeding seven times sd from background contraction, or 2% max force in the current trial (force task) or 5% max angle of the current trial (angle and velocity tasks). MEP amplitude and all continuous variables were standardized per participant by subtracting participant mean and dividing with sd. Standardization allowed us to center the priors around 0, with an sd = 1.

#### Testing MEP variance explained by the winning models

To evaluate how much MEP amplitude variance is accounted for by the winning models, we calculated the Bayesian R-squared for posterior distributions using the ‘bayes-R2’ function from the ‘brms’ package in R for the six winning models.

#### Additional effects of preceding trial, age, and sex

We tested whether extra variables had additional explanatory effect of MEP amplitude, including *preceding parameter level,* which is the level of the immediately preceding trial, *preceding parameter measure,* which is the parameter measure of the immediately preceding trial, *difference between current parameter level and preceding parameter level,* which is the level from the current trial minus level from the immediately preceding trial, *difference between current parameter measure and preceding parameter measure,* which is the parameter measure from the current trial minus the parameter measure from the immediately preceding trial, *sex,* and *age*. Further, for the velocity task, we tested whether max angle added explanatory power to the MEP amplitude, and likewise we tested whether mean velocity added explanatory power to the MEP amplitude in the angle task. We tested the effect of each variable by adding them to the winning model for each of the six conditions.

## Results

### Behavioral measurements

We examined the precision of parameter measures of movements. We investigated whether participants could produce different levels of force, angle, and velocity in the three parameter tasks without visual feedback. Figure 2 shows the distribution of parameter measures of force (Fig 2A-C), angle (Fig 2D-F), and velocity (Fig 2G-I) distributed by parameter level for all participants, standardized per participant.

**Figure 2:**
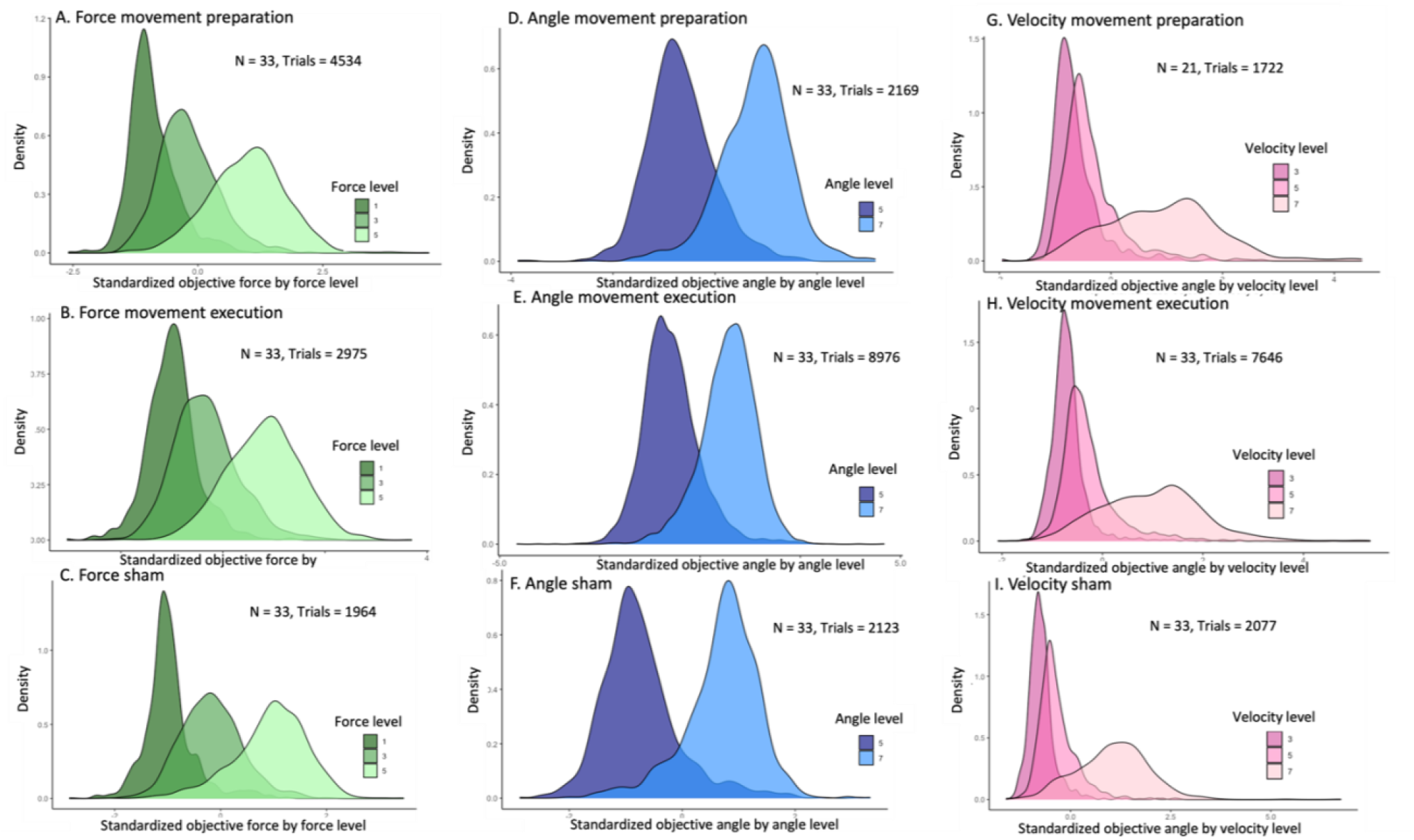
Measures of max force (A-C), max angle (D-F), and mean velocity (G-I), distributed by parameter level for all participants, standardized per participant.

#### Force: difference in distributions of force between force levels

Max force is distributed by force level 1, 3, and 5 (Fig. 2A-C). When estimating standardized max force as a function of subject and force level, the estimate increases for each force level across conditions, movement preparation, movement execution, and sham (Table 1). An estimate of 0 corresponds to the individual participant’s averaged max force across all trials. 95% CI intervals do not overlap between force levels in any condition and when tested with Bayesian hypothesis tests, all are different with infinite evidence ratio and posterior probability of 1 (Table 1). The width of 95% CI intervals for force level distributions do not differ between levels, indicating comparable precision for each force level. Taken together, participants can distinguish between the different force levels, and distributions are comparable to sham condition (Table 1).

**Table 1.** Distribution of parameter measures by parameter level standardized per participant. An estimate of 0 corresponds to the individual participant’s mean force, angle, or velocity across all trials. For all hypothesis tests across all conditions, evidence ratios are infinite and posterior probabilities are 1.

|  |  | Estimate | Est. Error | 95% CI | Hypothesis test | Estimate | Est. Error | 95% CI |
| --- | --- | --- | --- | --- | --- | --- | --- | --- |
| Force Prep. | Level 1 | -0.85 | 0.02 | [-0.89; -0.81] | Level 3 - level 1 > 0 | 1.56 | 0.04 | [1.50; 1.63] |
|  | Level 3 | 0.71 | 0.02 | [0.67; 0.76] | Level 5 - level 3 > 0 | 1.12 | 0.02 | [1.08; 1.15] |
|  | Level 5 | 1.83 | 0.02 | [1.78; 1.88] |  |  |  |  |
| Force Move | Level 1 | -1.04 | 0.03 | [-1.11; -0.98] | Level 3 - level 1 > 0 | 1.74 | 0.06 | [1.65; 1.84] |
|  | Level 3 | 0.70 | 0.03 | [0.63; 0.76] | Level 5 - level 3 > 0 | 1.12 | 0.03 | [1.07; 1.16] |
|  | Level 5 | 1.81 | 0.03 | [1.75; 1.88] |  |  |  |  |
| Force Sham | Level 1 | -1.0 | 0.03 | [-1.07; -0.93] | Level 3 - level 1 > 0 | 1.81 | 0.06 | [1.71; 1.91] |
|  | Level 3 | 0.81 | 0.03 | [0.74; 0.88] | Level 5 - level 3 > 0 | 1.15 | 0.03 | [1.10; 1.21] |
|  | Level 5 | 1.96 | 0.03 | [1.90; 2.03] |  |  |  |  |
| Angle Prep. | Level 5 | -0.69 | 0.03 | [-0.75; -0.64] | Level 7 - level 5 > 0 | 2.18 | 0.05 | [2.1; 2.26] |
|  | Level 7 | 1.49 | 0.03 | [1.43; 1.54] |  |  |  |  |
| Angle Move | Level 5 | -0.73 | 0.01 | [-0.75; -0.71] | Level 7 - level 5 > 0 | 2.16 | 0.02 | [2.12; 2.19] |
|  | Level 7 | 1.43 | 0.01 | [1.40; 1.46] |  |  |  |  |
| Angle Sham | Level 5 | -0.76 | 0.03 | [-0.81; -0.71] | Level 7 - level 5 > 0 | 2.25 | 0.05 | [2.17; 2.33] |
|  | Level 7 | 1.49 | 0.03 | [1.43; 1.54] |  |  |  |  |
| Velocity Prep. | Level 3 | -0.46 | 0.04 | [-0.53; -0.39] | Level 5 - level 3 > 0 | 0.73 | 0.06 | [0.62; 0.83] |
|  | Level 5 | 0.27 | 0.04 | [0.20; 0.34] | Level 7 - level 5 > 0 | 0.86 | 0.06 | [0.77; 0.95] |
|  | Level 7 | 1.13 | 0.06 | [1.01; 1.25] |  |  |  |  |
| Velocity Move | Level 3 | -0.51 | 0.02 | [-0.55; -0.48] | Level 5 - level 3 > 0 | 0.84 | 0.03 | [0.79; 0.89] |
|  | Level 5 | 0.33 | 0.02 | [0.30; 0.37] | Level 7 - level 5 > 0 | 0.95 | 0.02 | [0.91; 0.99] |
|  | Level 7 | 1.29 | 0.03 | [1.23; 1.34] |  |  |  |  |
| Velocity Sham | Level 3 | -0.63 | 0.03 | [-0.69; -0.57] | Level 5 - level 3 > 0 | 0.98 | 0.06 | [0.89; 1.08] |
|  | Level 5 | 0.36 | 0.03 | [0.30; 0.42] | Level 7 - level 5 > 0 | 1.21 | 0.04 | [1.14; 1.28] |
|  | Level 7 | 1.57 | 0.04 | [1.48; 1.66] |  |  |  |  |

#### Angle: difference in distributions of max angle between angle levels

Max angle is distributed by angle levels 5 and 7 (Fig. 2D-F). When estimating standardized max angle as a function of subject and angle level, the estimate of level 5 is smaller than level 7 across all conditions, movement preparation, movement execution and sham (Table 1). An estimate of 0 corresponds to the individual participant’s averaged max angle across all trials. 95% CI intervals do not overlap and when tested with Bayesian hypothesis testing, they are different with infinite evidence ratio and posterior probability of 1 (Table 1). The width of 95% CI intervals for angle level distributions do not differ substantially between levels, indicating comparable precision for each angle level. Taken together, participants can distinguish between angle levels, and distributions are comparable to sham condition (Table 1).

#### Velocity: difference in distributions of mean velocity between velocity levels

Mean velocity is distributed by velocity levels 3, 5, and 7 (Fig. 2h-j). When estimating standardized mean velocity as a function of subject and velocity level, the estimate increases for each level across all conditions, movement preparation, movement execution and sham. An estimate of 0 corresponds to the individual participant’s averaged mean velocity across all trials. As with force and angle, 95% CI intervals do not overlap and when tested with Bayesian hypothesis, they are different with infinite evidence ratio (Table 1). Taken together, participants can distinguish between velocity levels, and distributions are comparable to sham condition (Table 1). The width of 95% CI intervals for velocity level distributions differ between levels (Table 1). The increased distribution width for level 7 indicates that participants have greater difficulty in precision of high velocity levels compared with the lower velocity levels.

Taken together for all parameter measures (max force, max angle, and mean velocity), estimates increase with parameter level across all movement parameters for all conditions. Further, the evidence ratio (ER) for the difference between the distribution of parameter measures between parameter levels is infinite for all movement parameters across all conditions. Importantly, this demonstrates that the participants performed the tasks successfully and can distinguish between levels for all movement parameters based on internal monitoring of efferent and afferent signals. Taken together, participants can distinguish between the different levels of force, angle, and velocity.

#### Reaction Time

Mean reaction times for movement preparation are approximately 300ms (Table 2). In the movement preparation trials, the pre-movement stimulation delays movement onset, resulting in longer reaction times than for movement execution. Sham reaction times are comparable to movement execution reaction times. Time from TMS to movement onset in movement preparation is timed to 75ms after the go-tone, and participants initiate their movement approximately 200ms after the stimulation. TMS to movement onset times are negative for movement execution trials, since participants initiate the movement prior to TMS stimulation in these trials.

**Table 2.** Mean reaction times (ms, 1^st^ and 3^rd^ quartile) and mean time from TMS to movement onset (ms, 1^st^ and 3^rd^ quartile). Force preparation: Participants = 33, trials = 4534. Angle preparation: Participants = 33, trials = 2169. Velocity preparation: Participants = 21, trials = 1722. Force movement: Participants = 33, trials = 2975. Angle movement: Participants = 33, trials = 8976. Velocity movement: Participants = 33, trials = 7646. Force sham: Participants = 33, trials = 1964. Angle sham: Participants = 33, trials = 2123. Velocity sham: Participants = 33, 2077.

|  | RT (mean ms (1st Qu, 3rd Qu)) | Time from TMS to move onset (mean ms (1st Qu, 3rd Qu)) |
| --- | --- | --- |
| Force preparation | 329 (192,429) | 255 (118,355) |
| Angle preparation | 264 (146,304) | 181 (65,227) |
| Velocity preparation | 327 (182,341) | 239 (87,250) |
| Force movement | 139 (75,341) | -175 (-214,-1) |
| Angle movement | 163 (-8,306) | -164 (-165,-43) |
| Velocity movement | 316 (100,514) | -245 (-312,-52) |
| Force sham | 218 (184,367) |  |
| Angle sham | 177 (16,328) |  |
| Velocity sham | 321 (106,508) |  |

### Movement parameters reflected in MEP amplitude

We investigated whether force-, angle-, and/or velocity-measures have explanatory power for the MEP amplitude during movement preparation and movement execution. We estimated Bayesian linear mixed models using brms for all six conditions and compared them using bayes_factor in a forward feature selection framework (Table 3), see *methods* section for overview of included variables. The order of the variables in each model signifies the explanatory power of the variable. The first variable has most explanatory power. The winning model for each condition is seen in Table 4.

**Table 3.**
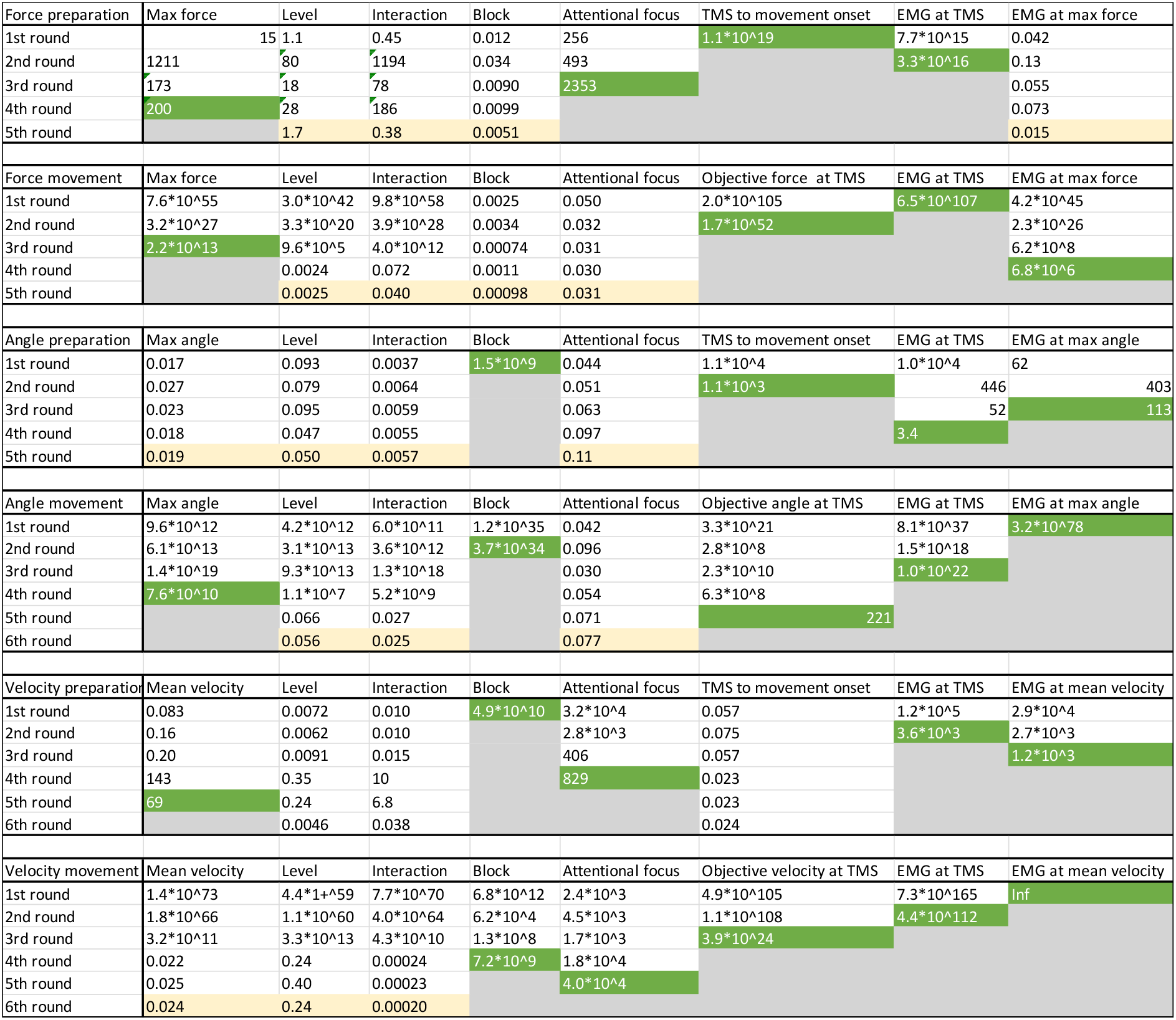
Forward feature selection of winning models tested with Bayesian model comparison for each parameter force, angle, and velocity during movement preparation and movement execution. Subject is included in all models before testing the other variables. For each selection round, the variable with largest bayes factor (BF) is included in the model (green). When no more variables have BF > 3, the winning model has been identified.

**Table 4.** Winning models of variables explaining MEP amplitude. For preparation models, TMS stimulation is induced before movement onset, and for execution models, TMS stimulation occurs during movement execution.

| Winning Models of Forward Feature Selection explaining MEP size |  |
| --- | --- |
| Force preparation | Subject + TMS to movement onset + EMG at TMS + attentional focus + max force |
| Angle preparation | Subject + block + TMS to movement onset + EMG at max angle + EMG at TMS |
| Velocity preparation | Subject + block + EMG at TMS + EMG at max velocity + attentional focus + mean velocity |
| Force execution | Subject + EMG at TMS + force at TMS + max force + EMG at max force |
| Angle execution | Subject + EMG at max angle + block + EMG at TMS + max angle + angle at TMS |
| Velocity execution | Subject + EMG at max velocity + EMG at TMS + velocity at TMS + block + attentional focus |

### Movement preparation

#### Force

For force with TMS stimulation during preparation, the variables *subject, time from TMS to movement ons*et (*BF* = 1.1 ∗ 10^19^), *EMG at TMS* (*BF* = 3.3 ∗ 10^16^), *attentional focus* (*BF* = 2353), and *max force* (*BF* = 200) are included in the model with most explanatory power of MEP. Importantly, *max force* is reflected in the MEP during movement preparation (slope = 0.063, evidence ratio = Inf) (Fig. 3 and Supplementary Table 1).

**Figure 3.**
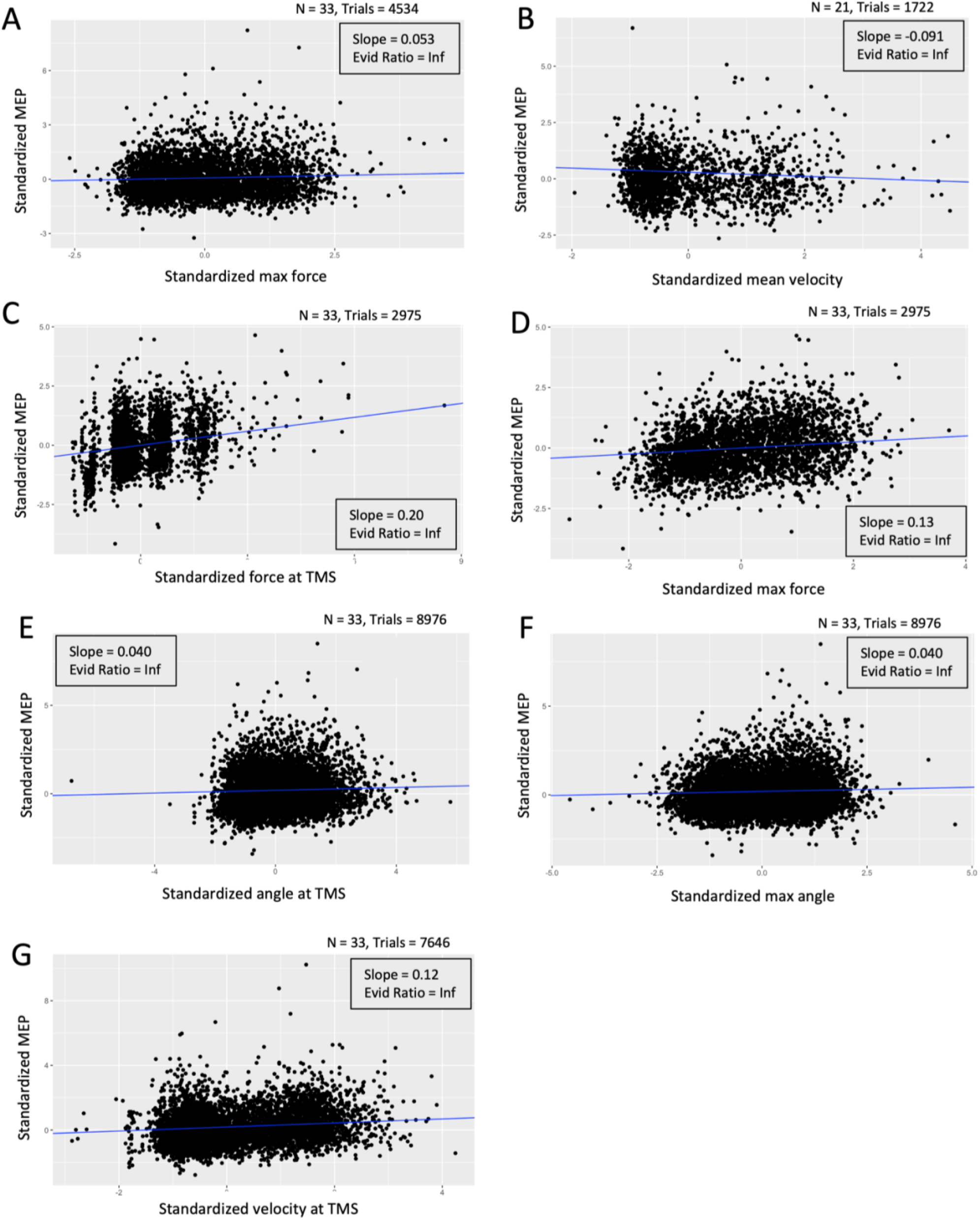
Correlation of standardized parameter measures with standardized MEP amplitude. A)-B) TMS stimulation during movement preparation. C)-G) TMS stimulation during movement execution. A) Standardized max force of ensuing trial by standardized MEP amplitude during movement preparation. B) Standardized mean velocity of ensuing trial by standardized MEP amplitude during movement preparation. C) Standardized force at TMS by standardized MEP amplitude during movement execution. D) Standardized max force by standardized MEP amplitude during movement execution. E) Standardized angle at TMS by standardized MEP amplitude during movement execution. F) Standardized max angle by standardized MEP amplitude during movement execution. G) Standardized velocity at TMS by standardized MEP during movement execution.

#### Angle

For angle with TMS stimulation during preparation, the variables *subject, block* (*BF* = 1.5 ∗ 10^9^), *time from TMS to movement onset* (*BF* = 1.1 ∗ 10^3^), *EMG at max angle* (*BF* = 113), and *EMG at TMS* (*BF* = 3.4) are included in the winning model. Importantly, *max angle* is not reflected in the MEP during movement preparation.

#### Velocity

For velocity with TMS stimulation during preparation, the variables *subject, block* (*BF* = 4.9 ∗ 10^10^), *EMG at TMS* (*BF* = 3.3 ∗ 10^3^), *EMG at max velocity* (*BF* = 1.2 ∗ 10^3^), *attentional focus* (*BF* = 829), and *mean velocity* (*BF* = 69) are included in the winning model. Importantly, *mean velocity* is reflected in the MEP amplitude during movement preparation (slope = -0.91, evidence ratio = Inf) (Fig. 3 and Supplementary Table 1).

Taken together, for force and velocity tasks, during preparation for movement both max force and mean velocity are reflected in the MEP. Similarly, for force and velocity tasks, attentional focus is included. Time from TMS to movement onset is explanatory for MEP during force and angle movement preparation.

### Movement execution

#### Force

For force with TMS stimulation during movement, the variables *subject, EMG at TMS* (*BF* = 6.5 ∗ 10^107^), *force at TMS* (*BF* = 1.7 ∗ 10^52^), *max force* (*BF* = 2.2 ∗ 10^13^), and *EMG at max force* (*BF* = 6.8 ∗ 10^6^) were included in the winning model. Importantly, both force-related measures, max force (slope = 0.13, evidence ratio = Inf) and force at TMS (slope = 0.20, evidence ratio = Inf) are included in the winning model (Fig. 3 and Supplementary Table 1).

#### Angle

For angle with TMS stimulation during movement, the variables *subject, EMG at max angle* (*BF* = 3.2 ∗ 10^78^), *block* (*BF* = 3.7 ∗ 10^34^), *EMG at TMS* (*BF* = 1.0 ∗ 10^22^), *max angle* (*BF* = 7.6 ∗ 10^10^), and *angle at TMS* (*BF* = 221) are included in the winning model. Importantly, both angle-related measures, max angle (slope = 0.040, evidence ratio = Inf) and angle at TMS (slope = 0.040, evidence ratio = Inf) are included in the winning model (Fig. 3 and Supplementary Table 1).

#### Velocity

For velocity with TMS stimulation during movement, the variables *subject, EMG at max velocity* (*BF* = *Inf*), *EMG at TMS* (*BF* = 4.4 ∗ 10^112^), *velocity at TMS* (*BF* = 3.9 ∗ 10^24^), *block* (*BF* = 7.2 ∗ 10^9^), and *attentional focus* (*BF* = 4.0 ∗ 10^4^) are included in the winning model. Importantly, *velocity at TMS* (slope = 0.12, evidence ratio = Inf) is included in the winning model, but not *mean velocity*.

Taken together, for movement execution, measures of force, angle, and velocity are all included in the winning models explaining the MEP.

### Additional effects of extra variables

We tested additional effects of extra variables mentioned in *methods.* For all variables tested as additional effects to the six winning models, only one variable added explanatory power. This was *last max force* (BF = 3.6) in force preparation.

### MEP amplitude modulation variance explained by winning models

Computing a BF provides an estimate of how much better one model is compared to another, but is not informative of the quality of the models. To test how much of the variance in MEP amplitude modulations the winning models explain, we estimated a Bayesian version of R-squared (Table 6). We found that for the movement preparation models, force R-squared was estimated to be: Est. = 0.0415 (Est. Error = 0.00538, Q2.5 = 0.0315, Q97.5 = 0.0526), angle R-squared Est. = 0.0453 (Est. Error = 0.00748, Q2.5 = 0.317, Q97.5 = 0.0607), and velocity R-squared Est. = 0.0789 (Est. Error = 0.0111, Q2.5 = 0.0576, Q97.5 = 0.101). For the movement execution models, force R-squared was estimated to be: Est. 0.225 (Est. Error = 0.111, Q2.5 = 0.203, Q97.5 = 0.101), angle R-squared Est. = 0.0584 (Est. Error = 0.00357, Q2.5 = 0.0516, Q97.5 = 0.0656), and velocity R-squared Est. = 0.268 (Est. Error = 0.00727, Q2.5 = 0.254, Q97.5 = 0.282). Estimates of MEP amplitude variance explained by the winning models are highest for velocity in both the movement preparation and movement execution condition. A high estimate is also found for variance explained by the force winning model in movement execution.

**Table 6.** R-squared estimates of the winning models.

|  | R2 estimate | Est. Error | Q2.5 & Q97.5 |
| --- | --- | --- | --- |
| Force preparation | 0.0415 | 0.00538 | [0.0315; 0.0526] |
| Angle preparation | 0.0453 | 0.00748 | [0.0317; 0.0607] |
| Velocity preparation | 0.0789 | 0.0111 | [0.0576; 0.101] |
| Force movement | 0.225 | 0.111 | [0.203; 0.247] |
| Angle movement | 0.0584 | 0.00357 | [0.0516; 0.0656] |
| Velocity movement | 0.268 | 0.00727 | [0.254; 0.282] |

## Discussion

The aim of this study was to investigate whether the movement parameters force, angle, and velocity, are reflected in the MEP amplitude to better understand the importance of these parameters in motor control. We found that during movement execution, all three movement parameters are reflected in the MEP amplitude modulations when correcting for EMG level. During movement preparation, velocity, and force, but not angle, are reflected in the MEP amplitude when correcting for EMG level. R-squared estimates of the winning models indicates that velocity and force models in movement execution and the velocity model in movement preparation explain a relevant part of the variance in MEP amplitude modulations. These results indicate that the CSE is modulated with all three movement parameters and suggests a flexibility in the control of movement parameters at the CSP level beyond the CSE modulation by EMG level.

### Behavior

On a group level, participants can distinguish between the instructed levels of all parameters without visual feedback, indicated by infinite evidence ratios of the differences in parameter measures between parameter levels. Regarding behavior, precision rather than accuracy has been the focus in this study, since without external feedback it is more relevant whether participant can consistently reproduce parameter levels rather than match externally defined levels. For force and angle, participants produce each level with equal precision, indicated by similar credibility interval (CI) widths. Prior research find a lower acuity for index finger velocity than for angle during passive movement (Long et al., 2022), which is supported by our results that show a more uneven distribution of measures of velocity compared to angle, and we further see a more precise distinction between force levels than velocity levels. In the velocity task, precision decreases with higher velocity, indicated by wider CI for level 7 than for level 3 and 5. Together, behavioral results confirm that participants perform the tasks successfully and can distinguish between force, angle, and velocity levels.

### Movement parameters reflected in MEP amplitude modulations

#### Movement execution

Through Bayesian model selection in a forward feature selection framework, we identified a winning linear mixed model for each parameter force, angle, and velocity during movement preparation and movement execution, to investigate whether the parameter measures of interest in each respective task were present in the model. In each winning model for movement execution, measures of movement parameters have explanatory power on MEP amplitude modulations. Measures of all three parameters are positively correlated with MEP amplitude during movement execution. Our results imply that the CSE can reflect either force, angle, or velocity of a movement, depending on the task that is performed. This suggest that the motor system is flexible in its information processing and highlights the capacity of the CSP to tune its excitability to varying task requirements.

#### Movement preparation

During movement preparation, the neural activity of the cortical motor system and the CSE changes in preparation for the movement (Bestmann and Duque, 2016; Cohen et al., 2010; Dupont-Hadwen et al., 2019). In our study, we find that force and velocity but not angle are reflected in the MEP amplitude prior to movement onset. This indicates a fine-tuning of the CSE to reflect information of task specific parameters of velocity and force already during movement preparation. Interestingly, *mean velocity* is negatively correlated with MEP amplitude modulations (slope = -0.092, evidence ratio (ER) = Inf), whereas force is positively correlated with MEP amplitude modulations during movement preparation (slope = 0.053, ER = Inf). The different directions of effect of force and velocity suggests that during movement preparation, the CSE is modulated independently in relation to force and velocity.

#### Parameter modulations

R-squared tests revealed that the winning models for force and velocity during movement execution explain a relevant degree of variance in the MEP amplitude modulations, and to a smaller degree the angle winning model. During movement preparation, the winning model for velocity explain more of the variance in the MEP amplitude modulations compared to models for force and angle. While we cannot definitively attribute this difference to the parameter measures, it would be a likely explanation since all other variables are similar between the models or had equal possibility to be included. Consequently, our findings suggest that CSE modulations are more closely related to velocity measures during movement preparation than to force and angle measures, and more closely related to velocity and force measures during movement execution than angle measures. This preference for velocity measures have prior been indicated for M1 activity (Stark et al., 2007).

Interestingly, velocity is negatively correlated with MEP amplitude modulations in the movement preparation condition, whereas it is positively correlated with MEP amplitude modulations in the movement execution condition. This indicates that these phases are neurophysiologically distinct. MEP amplitude is known to decrease during movement preparation and increase at movement onset (Bestmann and Duque, 2016). Research in pre-movement spinal activity in monkeys found that a majority of spinal neurons exhibited inhibitory activity during movement preparation and excitatory activity during movement execution (Prut and Fetz, 1999). Prut and Fetz also observed these opposite directed activation pattern in M1 (Prut and Fetz, 1999). The negative relation between velocity and MEP amplitude modulations we detect during movement preparation could therefore be attributed to increased inhibitory mechanisms in preparation for higher velocities. Subsequently, during the execution phase, the positive correlation between velocity and MEP amplitude indicates heightened excitation required to generate the higher velocity.

Although *max angle* is reflected in the MEP during movement execution, we do not see it reflected during movement preparation. The motor system needs proprioceptive and sensorimotor feedback for precise motor control (Schwartz, 2016). During movement preparation, the brain does not receive the same degree of sensory feedback as during movement execution. Integration of feedback with the outgoing signal allows the brain to compare the difference between current and intended position. The observed relation between MEP and *max angle* during movement execution as opposed to during movement preparation may therefore be attributed to a higher degree of sensorimotor integration of angle information with the motor command during movement execution than preparation.

Our findings demonstrate that modulations in MEP amplitude correlate with changes in movement parameters, however we cannot definitively assert that changes in MEP amplitude are the direct cause of these movement parameters. The observed changes in MEP amplitude reflect alterations in the physiological state of the CSP, which may result from either a causal relationship or a common underlying factor influencing both variables. Nevertheless, it suggests a functional linkage between the CSP activity and measures of movement parameters. Should there be a causal relationship, our results would imply that the advantages of being able to precisely control all parametric details of a movement outweigh the potential complexities introduced by increased degrees of freedom in the motor system. This enhanced control could indicate an evolutionary benefit in the fine-tuning of motor actions, crucial for complex index finger movements.

The variable attentional focus is included both velocity and force preparation and velocity movement execution models, which indicates differentiated modulation of CSE with different foci of attention during these conditions.

Changes in MEP amplitude vary with changes in M1 neuronal activity but are also affected by changes in afferent input to M1 from other cortical areas, peripheral somatosensory feedback, and by changes in spinal neuronal pool excitability (Bonnesen et al., 2022; Sasaki et al., 2018). Since MEP amplitude is influenced by both cortical and spinal contributions, the observed modulations could originate from either source. Further, the TMS-induced neural discharge pattern is not likely to exactly match the neuronal activation pattern elicited during voluntary movement (Bestmann and Krakauer, 2015). This could explain the variance in the MEP amplitude modulations that the winning models do not account for.

Force, angle, and velocity levels are most likely positively correlated. It could therefore be argued that one parameter potentially could be a confounding factor for another parameter, e.g. higher force levels would lead to higher velocity levels. Importantly, in movement preparation, the force estimate is positive (estimate = 0.05, estimated error = 0.01) whereas the velocity estimate is negative (estimate = -0.09, estimated error = 0.02).

This indicates different CSE modulation between the parameters since force positively modulates MEP amplitude and velocity negatively modulates MEP amplitude during movement preparation, allowing us to distinguish between the modulation of the different movement parameters.

### EMG is reflected in the MEP amplitude

The variable EMG at TMS is included in all six winning models, whereas the variable EMG at objective measure is included in all movement execution models and the angle and velocity movement preparation models. The parameter measures are represented in the models in addition to the EMG variables, indicating that the modulation of MEP amplitude by the three movement parameters are taking place independent of the EMG modulation of the MEP amplitude.

Contraction of the target muscle induces facilitation of the MEP amplitude (Hess et al., 1987). Muscle activity is therefore potentially a confounding factor, especially in the movement execution conditions. To control for this, we included the variable *EMG at time of TMS* in our analysis. As expected, it was through forward feature selection included as a variable in the winning model across all six conditions. When it is included, we ensure that if parameter measures are added to the model, it is because they provide additional explanatory power over and above EMG. To further isolate the relation between parameter measures and the MEP amplitude, we included the variable *EMG at parameter measure* in the analysis. This variable was through forward feature selection included in the winning model in five of six conditions, *force preparation* being the exception. Through the forward feature selection approach, the variable with most explanatory power is selected first, and the degree of variability in the MEP amplitude is accounted for by this variable, before testing the added explanatory power of the next variable. Therefore, the next variable is only added to the model, if it adds explanatory power in addition to the previously selected variables. This approach ensures that if parameter measures are added, they are only added in virtue of providing further explanatory power to the MEP amplitude. We therefore believe that the explanatory effect of parameter measures on MEP amplitude is not a consequence of muscle contraction. This is further supported by the fact that a linear relationship between movement parameters and muscle contraction cannot be assumed for angle and velocity. The lack of a direct linear relationship between EMG and measures of velocity and angle will enable the model to identify whether there is an additional effect of parameter measures. We further see that the MEP modulation differ between movement parameters, in that velocity negatively modulates MEP amplitude during movement preparation and positively modulates MEP amplitude during movement execution (Fig. 3 and Supplementary Table 1). We therefore believe that parameter measures contribute explanatory power in addition to EMG at parameter measure and EMG at time of TMS stimulation, emphasizing their additional effect on the MEP amplitude.

### Movement preparation and reaction time

It is reasonable to consider whether pre-movement stimulation occurred during movement preparation and not before cognitive preparation for the movement or during movement execution. Participants were instructed to react as fast as possible to the go-tone in the force and angle tasks. Based on prior studies, reaction time (RT) was expected to be approximately 150 ms (Leocani et al., 2000), and we observed comparable mean RTs in this study (force movement: 139 ms, angle movement: 163 ms). In movement preparation, we timed the TMS stimulation to occur 75 ms after initiation of the go-tone, for the stimulation to occur during the time the brain prepares the movement. Mean RT in movement preparation is for force 329 ms and angle 264 ms, which is longer than in the comparable movement execution trials. This delay is attributed to the disturbance caused by TMS stimulation during movement preparation, affecting the movement onset timing (Alibiglou and Mackinnon, 2012; Day et al., 1989), hence we are convinced that participants are in the movement preparatory phase at time of stimulation (Alibiglou and Mackinnon, 2012; Day et al., 1989). Participants were intentionally not instructed to react as fast as possible in the velocity task to avoid potential influence on mean velocity during the trial, resulting in similar RTs in all three conditions (preparation: 327 ms, movement: 316 ms, sham: 321 ms). Velocity preparation mean RT is comparable to that for force and angle, and we therefore believe that they are in the movement preparation phase at time of stimulation for velocity as well.

The MEP amplitude is modulated in preparation for the movement during movement preparation, making it important to consider the effect of time from stimulation to movement onset. The MEP is modulated in preparation for the movement non-linearly, exhibiting an initial decrease in MEP amplitude succeeding go-signal, followed by an increase in MEP amplitude up to movement execution (Bestmann and Duque, 2016; Derosiere et al., 2020). Davranche et al. (2007) found that an initial decrease in MEP area is proposed to be a cortico-spinal inhibition that ensures that cortical excitability can increase during action preparation, without premature movement initiation (Davranche et al., 2007). Due to this preparatory modulation of MEP amplitude, the timing of the TMS stimulation is an important determinant for the MEP amplitude in the movement preparation condition (Derosiere et al., 2020). To address this influence, we included a measure of time from TMS stimulation to movement onset as a variable in our analysis of movement preparation. As anticipated, the variable *time from TMS stimulation to movement onset* was included in the winning movement preparation model in both the force and angle task, but unexpectedly absent in the velocity movement preparation model. Whether this is because participants were not instructed to move right at the movement of the go-signal but were focusing on moving at a specific velocity may need to be further studied in detail.

The variable *block* has an explanatory effect on MEP amplitude in both movement preparation and execution for angle and velocity tasks. This could be due to mental or neuronal fatigue over time, as attention and arousal level potentially decrease over time. This variable is therefore included in the winning models to account for this effect.

### Limitations

The estimation of time from TMS stimulation to movement onset relies on measures of increased EMG and measures of change in movement parameters. Silent period and the MEP potentially disturb the estimation of the timing of movement onset since it is not possible to identify a change in EMG during this period (Hupfeld et al., 2020). Therefore, there is a risk that we overestimate the values of *time from TMS to movement onset*, which could be another reason this variable surprisingly is not included in the winning velocity movement preparation model. If movement onset is inaccurately estimated, the influence of the variable *time from TMS to movement onset* on the MEP may be underestimated. Further, potentially the MEP amplitude itself affects the variable *time from TMS to movement onset,* given the modulatory effect of MEP amplitude on silent period duration (Škarabot et al., 2019), hence possibly affecting movement onset identification. A second limitation to this study is that we do not measure all movement parameters in the same paradigm, which prevents us from evaluating their relative contribution to the MEP.

It is a limitation that EMG during MVC is only tested with isometric index finger movement, when we in the velocity and angle tasks obtain MEPs during flexion of index finger, since the muscle length will differ during flexion.

TMS induced perturbations of the movement, especially in the force task due to higher muscle activation. We evaluate that this is only a minor problem in the movement preparation condition, since pre-movement TMS delays movement rather than distorts it (Day et al., 1989). However, it might have introduced noise in the measures of movement parameters in the movement execution condition. The TMS stimulation occurred early during the movement, which would allow adjustments to realign with intended trajectories. However, the impact of these perturbations, alongside potential anticipatory effects on motor planning and sensory disturbances, cannot be discounted.

Data was collected over two 3-3.5 hour sessions separated by a long lunch-break. The duration of data collection introduced potential issues with physical and mental fatigue. Furthermore, continuous stimulation can potentially alter cortical excitability, which could affect MEP amplitudes (Ammann et al., 2020). To mitigate these effects, participants were encouraged to take regular breaks, and we ensured that they stayed hydrated and had access to snacks throughout the sessions. Despite these measures, our analysis reveals an impact of block on MEP amplitude in both angle and velocity models, although it is absent in the force models. The extension of the data collection provides a significant benefit by yielding large datasets.

### Conclusion, perspectives, and future studies

One important future extension of the present results would be an experimental setup in which all movement parameters can be modulated and measured simultaneously. This would allow us to examine the relative contribution of each movement parameter on explaining the MEP amplitude. It would also allow us to answer two questions. First, are the movement parameters reflected in the MEP amplitude all at once? Second, is the modulation of the MEP amplitude dependent on which movement parameter the participant is instructed to perform? This would allow us to explore whether the motor system reflects all movement parameters at the same time to accommodate the great complexity of human movement or whether it adjusts its output to the currently most important parameter to optimize the motor plan for task requirements.

The main finding of this study is that all movement parameters, force, angle, and velocity, are reflected in MEP amplitude modulations during movement, while force and velocity are reflected in MEP amplitude modulations during movement preparation, when correcting for EMG activity. This finding highlights a flexibility in the communication of the motor system, with distinct modulations depending on movement parameter in movement preparation, indicating differentiable preparatory mechanisms for the three movement parameters. Our findings suggest that the motor system uses an adaptive form of signaling, effectively reflecting information regarding all three parameters force, angle, and velocity.

## Additional information

### Data availability

We will make anonymized data and scripts available at Open Science Foundation upon acceptance of the manuscript. Data is currently pseudo-anonymized, so if reviewers suggest further processing of the data that may require any information regarding research participants, we cannot delete the pseudo-anonymization code and make the data fully anonymized. Therefore, data can only for legal reasons be made available upon manuscript acceptance with all suggested changes to data processing.

### Author contributions

Ida Marie Brandt: Conceptualized and designed the study, data collection, data analysis, discussed results. Writing: original draft.

Jesper Lundbye-Jensen: Conceptualized the study, provided feedback on experimental setup. Writing: Review and editing.

Thor Grünbaum: Conceptualized the study, provided feedback on experimental setup, discussed results. Writing: Review and editing.

Mark Schram Christensen: Obtained funding, conceptualized and designed the study, supervised data analysis, discussed results. Writing: Review and editing.

## Acknowledgements

The authors would like to thank Mark Wulf Carstensen and Iva Svecová for helping during data collection, Anke Ninija Karabanov and Victor Lange for theoretical contributions to experimental setup and Valeria Simonelli and Iva Svecová for contributing feedback during piloting.

## Supplementary data

**Figure 1.**
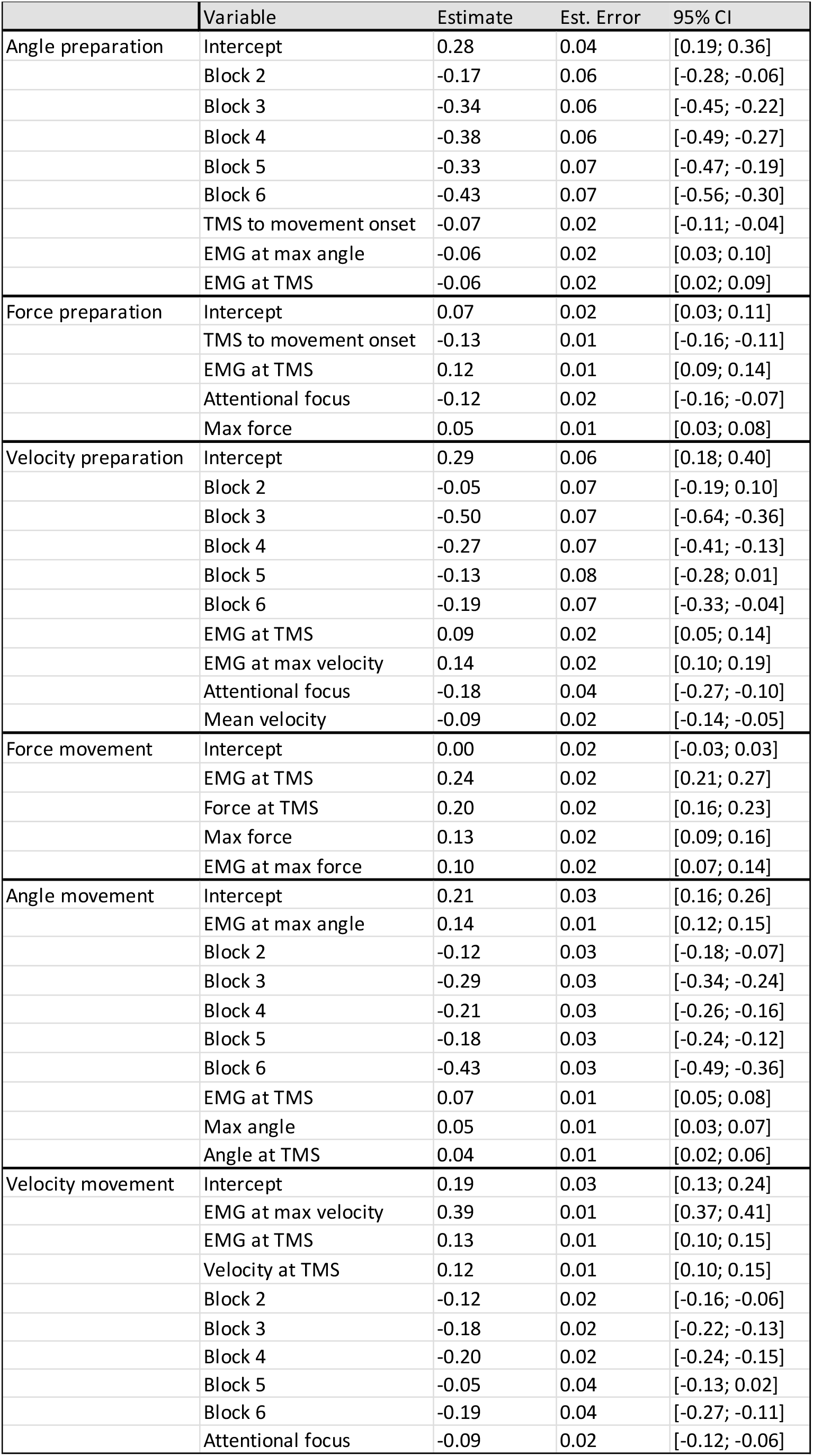
Bayesian linear mixed model estimates from the winning models for each condition. Parameter measures and MEP amplitudes are standardized.

